# Thymine DNA glycosylase combines sliding, hopping, and nucleosome interactions to efficiently search for 5-formylcytosine

**DOI:** 10.1101/2023.10.04.560925

**Authors:** Brittani L. Schnable, Matthew A. Schaich, Vera Roginskaya, Liam P. Leary, Tyler M. Weaver, Bret D. Freudenthal, Alexander C. Drohat, Bennett Van Houten

**Affiliations:** Molecular Biophysics and Structural Biology Graduate Program, University of Pittsburg, PA 15260, USA; UPMC Hillman Cancer Center, University of Pittsburgh, Pittsburgh, PA 15213, USA; Department of Pharmacology and Chemical Biology, School of Medicine, University of Pittsburgh, Pittsburgh, PA 15213, USA; Department of Biochemistry and Molecular Biology, Department of Cancer Biology, University of Kansas Medical Center, Kansas City, KS 66160, USA; Department of Biochemistry and Molecular Biology, University of Maryland School of Medicine, Baltimore, Maryland 21201, USA

## Abstract

Base excision repair is the main pathway involved in active DNA demethylation. 5-formylctyosine and 5-carboxylcytosine, two oxidized moieties of methylated cytosine, are recognized and removed by thymine DNA glycosylase (TDG) to generate an abasic site. Using single molecule fluorescence experiments, we studied TDG in the presence and absence of 5-formylctyosine. TDG exhibits multiple modes of linear diffusion, including hopping and sliding, in search of a lesion. We probed TDG active site variants and truncated N-terminus revealing how these variants alter the lesion search and recognition mechanism of TDG. On DNA containing an undamaged nucleosome, TDG was found to either bypass, colocalize with, or encounter but not bypass the nucleosome. However, truncating the N-terminus reduced the number of interactions with the nucleosome. Our findings provide unprecedented mechanistic insights into how TDG searches for DNA lesions in chromatin.

## Introduction

DNA can be damaged by a variety of endogenous and exogenous sources and these potentially mutagenic bases are removed via base excision repair (BER)^1^. In humans, BER is initiated by one of eleven damage specific DNA glycosylases that recognize a lesion and cleave the glycosidic bond to leave an abasic site. The repair is completed by the actions of several proteins that restore the correct base pair. Thymine DNA glycosylase (TDG) is a monofunctional glycosylase that initiates BER by recognizing a specific repertoire of lesions, including 5-methylcytosine (5mC) that has been deaminated to form a G:T mismatch or 5mC that has been oxidized by the ten eleven translocation (TET) family of enzymes into 5-formylcytosine (5fC) and 5-carboxylcytosine (5caC)^2–5^. Interestingly, TDG is the only known mammalian DNA glycosylase that creates an embryonic lethal phenotype in mice upon depletion: presumably due to the significant role that TDG plays in active demethylation, in addition to the action on environmental DNA damage^6,7^.

While TDG is known to preferentially remove T from G:T mismatches in the context of a CpG, other substrates do not show this sequence context dependence ^8,9^. One such lesion is 5fC, which has an abundance of 6×10^4^ per human genome, and the removal of 5fC was similar in the context of a CpG and outside a CpG^10–12^. Previous studies have shown that N140 of TDG is essential for the catalytic activity on 5fC because of interactions with the water nucleophile.

The N140A variant greatly reduces the catalytic activity 16,000-fold^5^. It has been shown that R275 plays an essential role in catalysis by filling in the void in the DNA that is generated once the substrate nucleobase is flipped into the active site^13^. In addition to TDG specificity for G:T mismatches and 5fC, TDG is able to bind to DNA nonspecifically with an affinity of 293 ± 64 nM and has a footprint of 10 base pairs^14,15^. While the roles for these key active site amino acids in catalysis is well-established, their role in mediating TDG searching for DNA lesions in the presence of excess unmodified DNA is unknown.

A major unexplored consideration for any glycosylase search process in a cell is the presence of chromatin. DNA packaged into nucleosomes would interrupt a DNA sliding mechanism and inhibit the accessibility of specific sites in the absence of nucleosome sliding and/or remodeling. Among DNA glycosylases this phenomenon is particularly relevant to TDG, as the methylated DNA where its substrates are formed are associated with higher degree of chromatinization^16^. TDG has also been shown to directly interact with nucleosome core particles, however kinetic studies also demonstrated TDG activity was only reduced ∼two-fold for uracil substrates present on a nucleosome core particle^17,18^.

Here, we utilized single-molecule analysis of DNA-binding proteins from nuclear extracts (SMADNE)^19^ to directly observe the search process of fluorescently labeled TDG in real-time on DNA containing 5fC substrates or nucleosome core particles. By observing TDG scanning the DNA as it searches for 5fC, we determined that TDG searches for damaged sites using a combination of sliding and hopping and that TDG diffusivity was highly dependent on DNA tension. This combined search mechanism was also observed on DNA containing NCPs. In some instances, TDG hopped over nucleosomes to continue searching the DNA, but in other cases it approached the NCP but did not bypass. We have also found that specific amino acid side chains that are essential for catalytic activity also impact the dwell time and the ability of TDG to diffuse on DNA. Additionally, although ΔN-term TDG (residues 82-410) is catalytically active, we demonstrate that the N-terminus plays a significant role in binding and searching unmodified DNA as well as playing a key role in engaging undamaged NCPs.

## Results

### TDG binds DNA specifically and nonspecifically

TDG-HaloTag was overexpressed in U2OS cells, and the nuclear extracts were quantified using western blots and SDS gels to determine a 60-fold difference in overexpressed fluorescently tagged WT TDG to endogenous TDG (Fig. S2). To determine the binding behavior of TDG on DNA containing a lesion of interest, we initially performed the single-molecule imaging with TDG-HaloTag-JF635 and DNA containing 5fC generated by nick translation (Fig. 1a, c). Analysis of the resulting kymographs revealed complex binding dynamics, including non-motile (12.6%) and motile TDG binding events (i.e., exhibited 1D diffusion, 87.4%). Further co-localization analysis of red TDG-HaloTag-JF635 with the blue 5fC fiducial marker in the DNA (fluorescein dUTP) revealed the stationary events were primarily TDG bound to 5fC sites (Fig. 1d). The motile events primarily represent TDG binding to the DNA nonspecifically. Of note, diffusion events that are less than 0.3 seconds are not detected. Fitting the motile events to a cumulative residence time distribution plot (CRTD) with a one-phase exponential decay, a binding lifetime of 7.5 ± 0.3 seconds was obtained (Fig. 1d-e, Table 1). Only motile events with a binding lifetime of 7.9 ± 0.1 sec were observed in additional single-molecule imaging experiments with TDG-HaloTag-JF635 and unmodified DNA, supporting that the stationary events reflect TDG bound to a lesion (Fig. 1f-g, Table 1). In contrast to the motile events, the CRTD plot of stationary events that co-localized with the 5fC fiducial marker were fit with a two-phase exponential decay with binding lifetimes of 14.7 ± 1.7 and 101.1 ± 7.1 seconds (Fig. 1d-e, Table 1). We hypothesize that these longer TDG lifetimes represent binding to and cleaving the 5fC. This is consistent with previous kinetic experiments that indicate it takes ∼ 68 secs (*k_max_* of 0.61 min^-1^) to cleave 5fC at 22°C, plus the time for TDG to dissociate from the generated abasic site at^2^. Notably, the shorter 14.7 ± 1.7 second binding events, that co-localized with the 5fC fiducial marker, exclusively occur after a long-lived event. These shorter events likely represent TDG binding to an abasic site generated by its catalytic function or a nicked abasic site processed by apurinic/apyrimidinic endonuclease 1 (APE1) in the extracts^19^.

**Figure 1:**
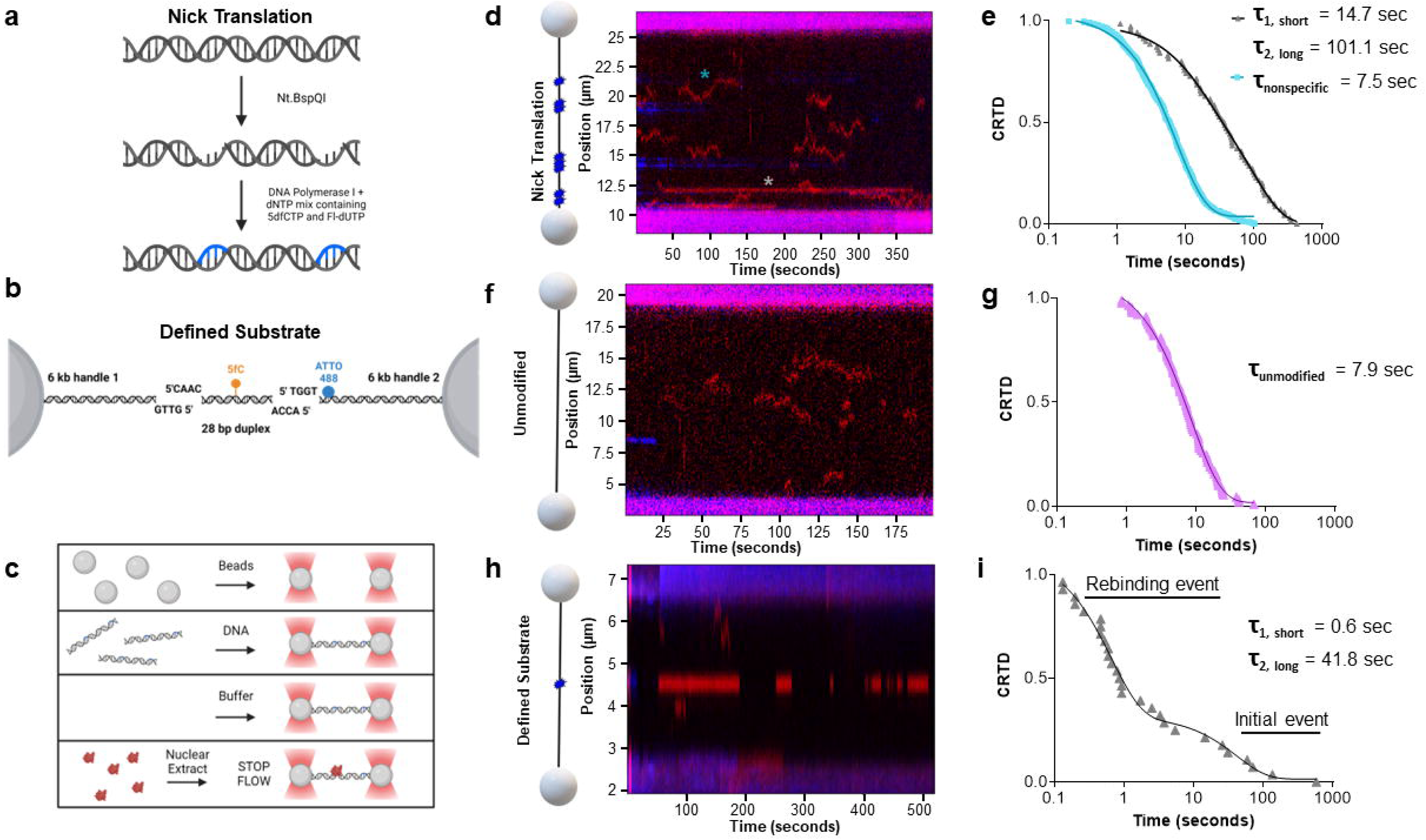
TDG binds DNA specifically and nonspecifically. (A) Cartoon schematic showing how 5fC was incorporated into nick translated λ DNA. (B) Cartoon schematic showing 28 base pair duplex DNA containing a single 5fC (orange) ligated into 6 kb LUMICKS handle kit, with handle 2 containing ATTO 488 (blue). (C) A diagram depicting the order of reagents, which are under laminar flow, are captured in the flowcell. (D) A cartoon depiction of the DNA substrate used for TDG binding, with 5fC sites shown in blue, and an example kymograph with TDG binding shown in red. Specific event indicated with grey asterisk and nonspecific event indicated with teal asterisk. (E) Cumulative Resident Time Distribution (CRTD) analysis fit to a two-phase decay of TDG binding DNA containing 5fC specifically (n = 70) and nonspecifically (n = 487). (F) A cartoon depiction of the unmodified DNA substrate with an example kymograph of TDG binding and moving. (G) CRTD analysis fit to a one-phase decay of TDG binding unmodified λ DNA (n = 155). (H) An example kymograph of TDG binding to a single 5fC. (I) CRTD analysis fit to a two-phase decay of TDG binding to 5fC (n = 28). Created with BioRender.com.

**Table 1.**
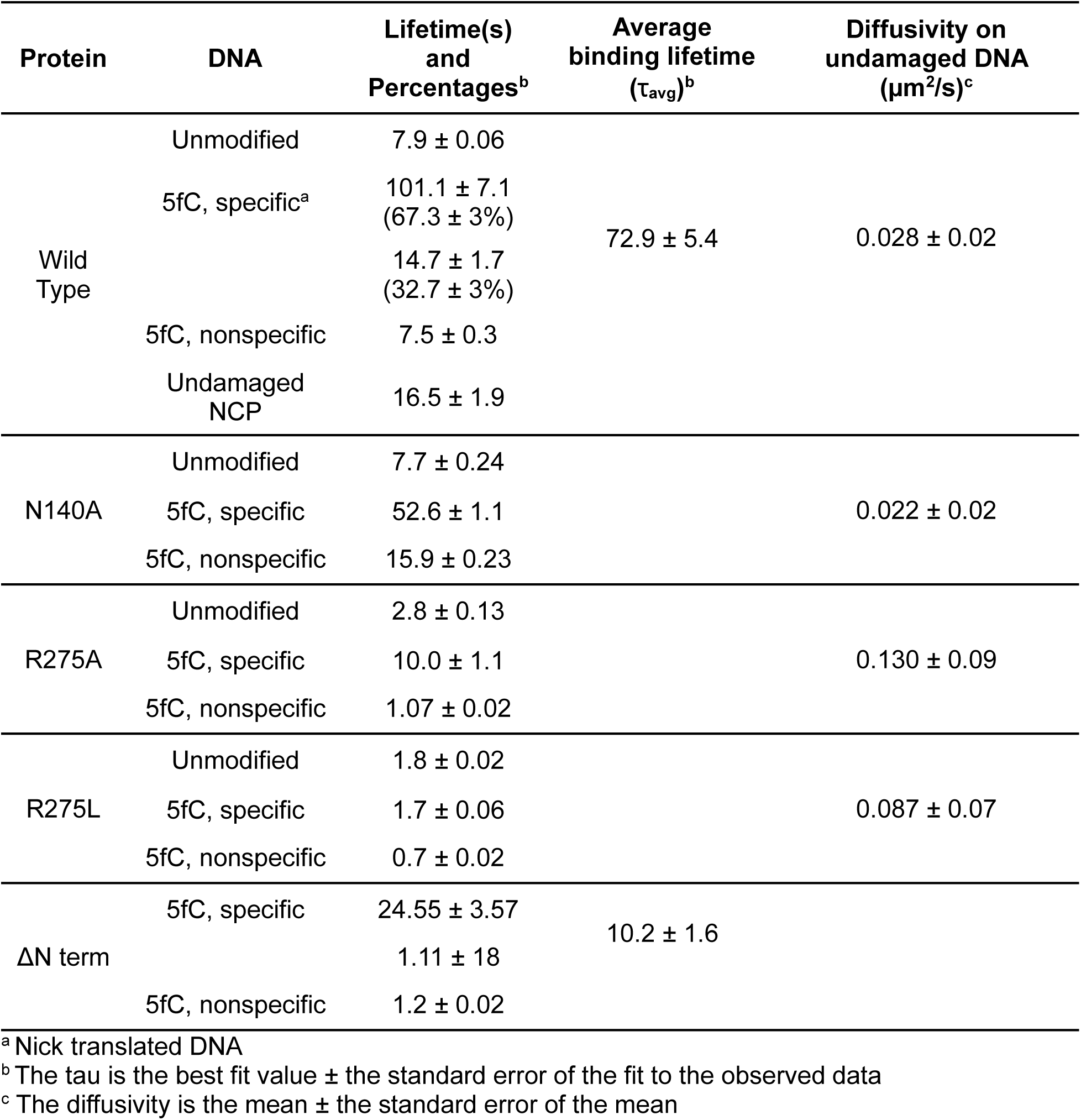
Binding lifetimes and diffusivity.

Since the nick translated DNA contains multiple sites with a region of 5fC, we generated a 12 kb DNA substrate that contains a single center-positioned 5fC to more easily observe if lifetime differs over time, indicating turnover. We then performed the correlative optical tweezers-fluorescence microscopy (CTFM) experiment using TDG-HaloTag-JF635 with the single 5fC DNA. A CRTD plot of stationary events that co-localized with the 5fC were fit with a two-phase exponential decay with binding lifetimes of 0.67± 0.06 sec and 41.8 ± 12.2 sec (Fig. 1d-e,h-i, Table 1), consistent with the short- and long-lived binding lifetimes observed on the DNA substrate containing 10 5fC sites (Fig. 1 e). These data further indicate that long-lived events represent TDG binding to the substrate and the short-lived events are mostly likely TDG rebinding to the catalytic product abasic site (Fig. 1h-i).

### TDG exhibits mixed modes of linear diffusion during its search for DNA damage

After observing TDG bind and move on DNA nonspecifically, we sought to determine the mode of linear diffusion TDG uses on DNA. Linear diffusion occurs through sliding and hopping mechanisms that have distinct behaviors. Sliding is when a protein tracks the DNA helix without dissociating, leading to coupled translational and rotational diffusion^20,21^. Alternatively, hopping is when a protein micro-dissociates as it translocates along the DNA, leading to uncoupled translational and rotational diffusions. Notably, the sliding and hopping mechanisms can be distinguished by determining whether the diffusion coefficient of TDG changes as a function of salt concentration: diffusion via sliding is unaffected by salt concentration, whereas diffusion for hopping is sensitive to salt concentration^22–24^. To further differentiate whether TDG uses a sliding, hopping, or multimodal 1D diffusion, we performed additional single-molecule imaging experiments with TDG-HaloTag-JF635 and unmodified DNA at varying ionic concentrations (75 to 150 mM NaCl). We observed an increase in the diffusion from 1.6×10^-2^ µm^2^/s at 75 mM NaCl, to 2.8×10^-2^ µm^2^/s at 100 mM NaCl, and 3.1×10^-2^ µm^2^/s at 150 mM NaCl (Fig. 2a, Table 1, Supplemental Fig. 5a-d). The theoretical limit of diffusion of TDG-HaloTag sliding along the DNA was calculated to be 2.56×10^-1^ µm^2^/s (see Supplemental Note). Any rates that exceed this theoretical limit of diffusion would reflect hopping events. While there was an increase in the diffusion with increasing ionic strength and a widening of the distribution of diffusivities, none of the values were above the theoretical limit of diffusion to unambiguously signify hopping. Since the crystal structures (PDBs 2RBA and 4Z7B) show TDG interacts with both strands of the helix when bound to a lesion, we labeled and mixed TDG-HaloTag with two different colored labels, JF635 (red) and JF552 (green) to directly visualize hopping. Two types of behavior were observed: (1) JF635-TDG and JF552-TDG collide, in which the two orthogonally labeled TDG proteins did not bypass one another (62.5%), and (2) JF635-TDG and JF552-TDG molecules bypass one another (37.5%) (Fig. 2b). The only way it is possible to observe both of these events is for TDG to have the ability to not only slide on the DNA but also to hop.

**Figure 2:**
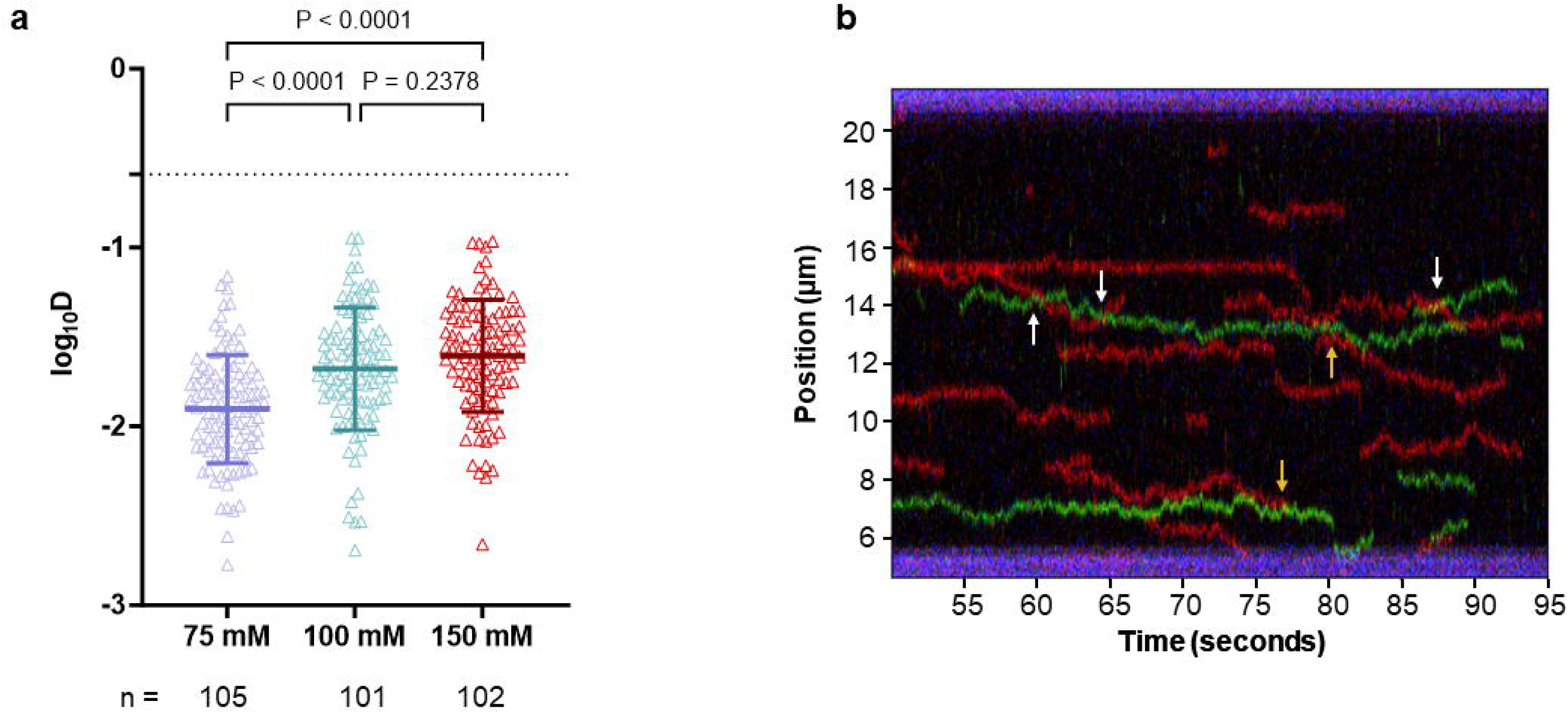
TDG exhibits linear diffusion on DNA. (A) Scatter plot of the diffusion coefficient (log_10_*D*) calculated for TDG with increasing ionic strength on unmodified λ. Dashed line, *D_lim_*, theoretical limit to free diffusion for TDG-HaloTag. ****p < 0.0001 by two-way ANOVA. (B) An example kymograph of the two-color TDG experiment. Separate TDG-HaloTag extracts were labeled with either JF635 (red) or JF552 (green) and mixed at a 1:1 ratio. White arrows indicate bypass events and yellow arrows indicate collision events.

### TDG catalytic variants reveal R275 is essential for base detection

N140 is essential for activating the water molecule for catalysis to occur and R275 fills in the hole in the duplex that is generated when the base is flipped into the active site. The difference in expression of each variant compared to endogenous TDG is 72-fold for N140A, 45-fold for R275A, and 21-fold for R275L (Fig. S2). Mutating to R275A and R275L decreases the excision of T from G:T pairs by 8-fold and 30-fold, respectively, and also reduces the binding affinity of either variant to G:T containing substrates 3-fold^13^. It is unknown if either variant affects the ability of TDG to search for DNA lesions^3,13,25^. To investigate the role of these key amino acids on the lesion search and recognition mechanism of TDG, we performed additional single-molecule imaging with TDG-HaloTag-N140A, R275L, and R275A variants and DNA containing regions of 5fC. Consistent with the WT TDG, we observed motile and non-motile events for all three variants tested, which represent nonspecific and 5fC binding, respectively. The N140A nonspecific events (motile) bound with a lifetime of 15.9 ± 0.23 seconds (84.8%) and the specific events (non-motile) bound with a lifetime of 52.9 ± 1.1 seconds (15.2%) (Fig. 3a, Supplemental Figs. 1d and 6a-b). This shorter lifetime for specific events (compared to WT) and the absence of short-lived rebinding events, further suggesting that wild type TDG is able to cleave the 5fC moiety and rebind to the abasic site product. The R275A nonspecific events (motile) bound with a lifetime of 1.07 ± 0.02 seconds (92.5%) and the specific events (non-motile) bound with a lifetime of 10.0 ± 1.1 seconds (7.5%) (Fig. 3b, Supplemental Figs. 1e and 6c-d). The R275L nonspecific events (motile) bound with a lifetime of 0.7 ± 0.02 seconds (88.7%) and the specific events (non-motile) bound with a lifetime of 1.7 ± 0.06 seconds (11.3%) (Fig. 3c, Supplemental Figs. 1e and 6e-f). N140A, R275A, and R275L bound to unmodified DNA with lifetimes of 7.7 ± 0.24, 2.8 ± 0.13, and 1.8 ± 0.02 seconds, respectively (Fig. 3a-c, Table 1), with the R275A and R275L lifetimes shorter than the lifetime of TDG-HaloTag-JF635 binding to unmodified DNA. Together these data reveal that disrupting key active site residues disrupts the ability of TDG to bind DNA as well as demonstrating a reduced binding lifetime at sites of 5fC, in agreement with previous bulk biochemical studies^13,26^.

**Figure 3:**
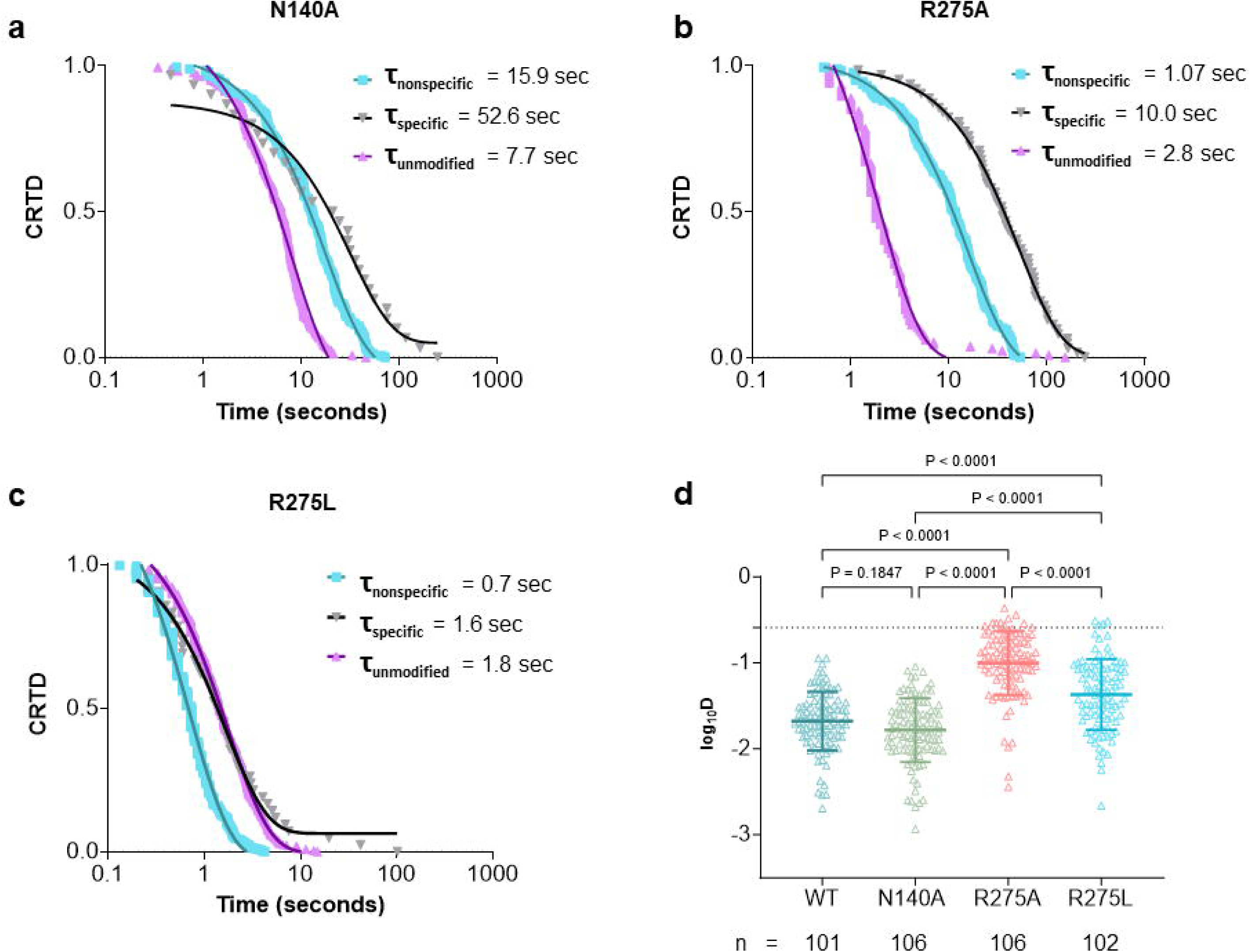
TDG catalytic variants indicate R275 is essential for base detection. Experiments conducted with unmodified or nick translated DNA containing 5fC. (A) CRTD analysis fit to a one-phase decay of N140A TDG binding to unmodified (n = 163), specifically to 5fC (n = 30) and nonspecifically (n = 168). (B) CRTD analysis fit to a one-phase decay of R275A TDG binding to unmodified (n = 110), specifically to 5fC (n = 24) and nonspecifically (n = 296). (C) CRTD analysis fit to a one-phase decay of R275L TDG binding to unmodified (n = 269), specifically to 5fC (n = 42) and nonspecifically (n = 330). (D) Scatter plot of the diffusion coefficient (log_10_*D*) calculated for each variant TDG on unmodified λ. Dashed line, *D_lim_*, theoretical limit to free diffusion for TDG-HaloTag. ****p < 0.0001 by two-way ANOVA.

Since we observed the binding lifetimes of the TDG variants to differ from wild type TDG, we examined the diffusion of these variants by mean squared displacement (MSD) analysis. MSD analysis measures the displacement of the particle over a given time window, over the course of the event length to determine the diffusivity and α, the anomalous diffusion coefficient. N140A had a diffusion coefficient of 2.2×10^-2^ µm^2^/s, which was not significantly different from wild type (Fig. 3d). This indicates that N140 does not significantly affect the movement of TDG on unmodified DNA, but the decreased binding lifetime leads to less DNA being sampled before dissociation. The diffusion coefficient for R275A (1.3×10^-1^ µm^2^/s) and R275L (8.7×10^-2^ µm^2^/s) were significantly faster than both the wild type and N140A variant (Fig. 3a, Table 1), suggesting that R275 is more important for searching unmodified DNA than N140.

We can estimate the residence time per base pair by utilizing the stepping rate (see Supplemental Note). The base pair residence time for wild type was estimated to be 2.1 ± 1.6 µs. For N140A, the base pair residence time was estimated to be 2.6 ± 2.2 µs, which is similar to the time for wild type TDG (2.1 ± 1.6 µs). In contrast, the R275A and R275L TDG variants had shorter residence times of 0.5 ± 0.3 µs and 0.7 ± 0.5 µs, respectively (Supplemental Table 1). This may indicate that the R275A/L variants are no longer able to flip out and sample base pairs, which agrees with previous biochemical studies that indicated R275 plays a role in promoting or stabilizing the flipped nucleotide^13^. While the faster diffusion observed for the R275A and R275L TDG variants increases the amount of DNA sampled during each binding event, the rapid search may be less effective if TDG is more likely to miss the lesion or modified base, similar to behavior observed with other DNA glycosylases wedge residue variants^27^.

### The N-terminus of TDG facilitates its movement on DNA

Previous work identified the first 81 amino acids are not essential for the catalytic activity of TDG^28^. However, we hypothesized that the intrinsically disordered N-terminus may play a key role for TDG search of unmodified DNA (Supplemental Fig. 1a, c). To investigate the role of the TDG N-terminal region on lesion search and lesion recognition mechanism, we performed additional single-molecule imaging experiments with TDG-HaloTag containing a truncation of the N-terminal 81 residues (ΔN-term-TDG-HaloTag) and DNA containing regions of 5fC modification. The ΔN-term-TDG-HaloTag was expressed 50-fold higher compared to endogenous TDG (Fig S2). We observed similar behavior with ΔN-term-TDG-HaloTag as full-length TDG-HaloTag, where long-lived, stationary events were followed by shorter lived stationary events (Fig. 4a). Interestingly, the nonspecific lifetime of ΔN-term-TDG-HaloTag was 1.2 ± 0.02 seconds which is three-fold shorter than full-length TDG-HaloTag (Fig. 4b). This reduced lifetime on nonspecific DNA supports the hypothesis that the N-terminus of TDG is important for searching the DNA.

**Figure 4:**
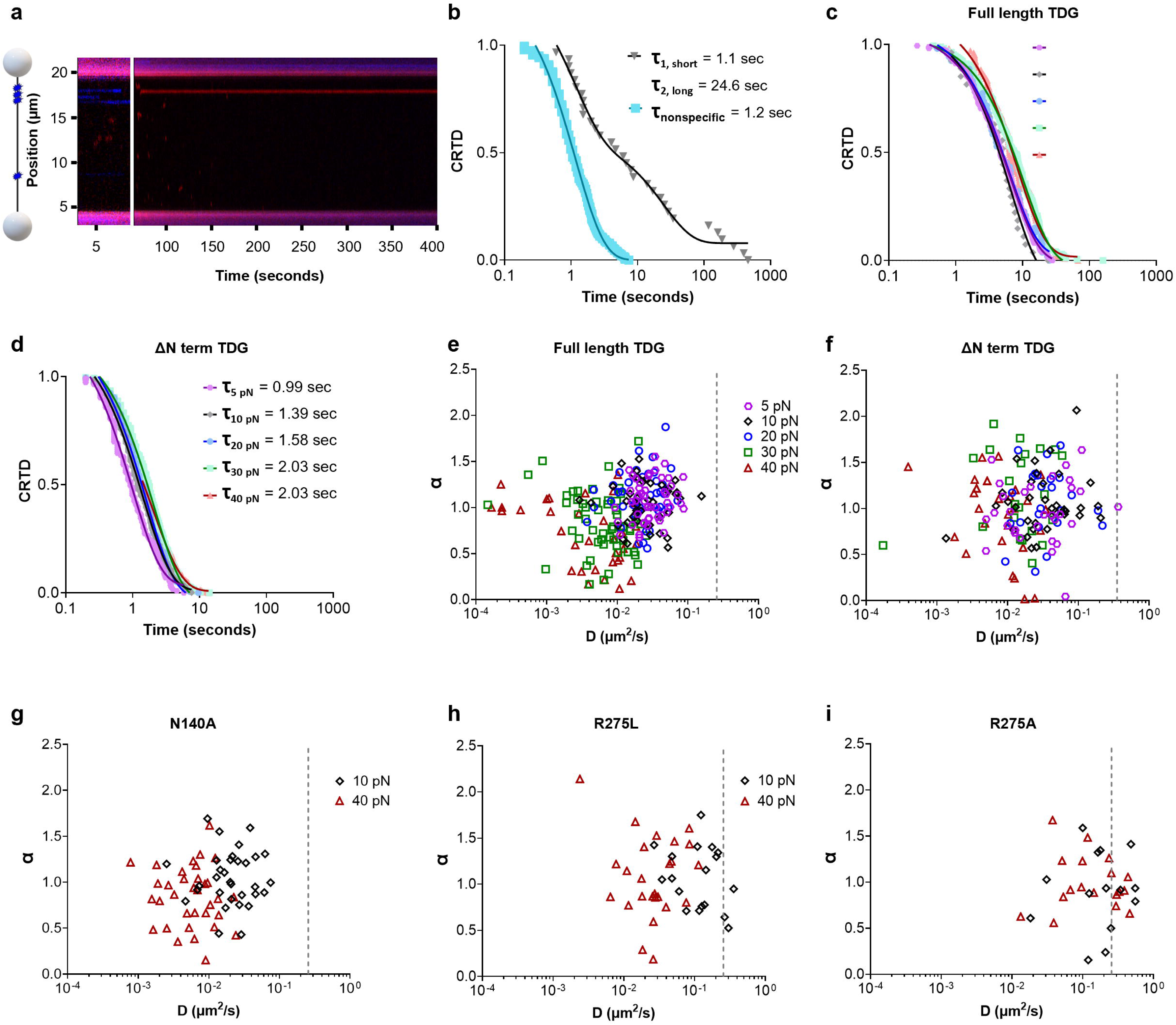
The N-terminus of TDG is significant for its movement on DNA. Tension alters diffusivity and α for ΔN-term and full-length TDG. (a) A cartoon depiction of the DNA substrate with 5fC sites shown in blue, and a representative kymograph with TDG binding shown in red. Break in kymograph to show location of fiducial markers. (b) CRTD analysis fit to a two-phase decay of TDG binding DNA containing 5fC specifically (n = 32). CRTD analysis fit to a one-phase decay of TDG binding nonspecifically (n = 243). (c) CRTD analysis fit to a one-phase decay of full-length TDG binding to unmodified DNA at 5 pN (n = 172), 10 pN (n = 27), 20 pN (n = 53), 30 pN (n = 169), and 40 pN (n = 102). (d) CRTD analysis fit to a one-phase decay of ΔN-term TDG binding to unmodified DNA at 5 pN (n = 113), 10 pN (n = 201), 20 pN (n = 197), 30 pN (n = 293), and 40 pN (n = 203) (e) Plot of diffusion coefficients (*D*) versus alpha (α) for full-length for 5 pN (n = 50), 10 pN (n = 41), 20 pN (n = 41), 30 pN (n = 57), and 40 pN (n = 33). (f) Plot of diffusion coefficients (*D*) versus alpha (α) for ΔN-term for 5 pN (n = 30), 10 pN (n = 36), 20 pN (n = 24), 30 pN (n = 22), and 40 pN (n = 35). (g) Plot of diffusion coefficients (*D*) versus alpha (α) for N140A for 10 pN (n = 33) and 40 pN (n = 34). (h) Plot of diffusion coefficients (*D*) versus alpha for 10 pN (n = 19) and 40 pN (n = 27). (α) for R275L. (i) Plot of diffusion coefficients (*D*) versus alpha (α) for R275A for 10 pN (n = 14) and 40 pN (n = 19).

A previous AFM study determined that TDG bends the DNA ∼30° when searching for a lesion, whereas the DNA is bent ∼70° when TDG is bound to the lesion^29^. We hypothesized based on the published AFM study^29^ that the necessity of DNA bending by TDG may result in shorter lifetimes at higher DNA tensions because TDG cannot fully engage with the extended DNA, but our data showed this is not the case. In our prior experiments we used a tension of 10 pN. By increasing the tension from 5 pN to 40 pN with full-length TDG, we observed similar lifetimes of 6.49 ± 0.02, 6.45 ± 0.06, and 6.25 ± 0.20 seconds at 5, 10 and 20 pN, respectively (Fig. 4c, Table 2). At 30 pN and 40 pN, we observed lifetimes that were significantly ∼two-fold higher than at lower tensions (Fig. 4c, Table 2). For ΔN-term TDG, we also observed a two-fold increase in the binding lifetimes with increasing tension. Additionally, ΔN-term TDG stays bound to the unmodified DNA with a 3-5-fold shorter lifetime than full-length at the respective tensions (Table 2). Shorter lifetimes for ΔN-term compared to full-length further supports that idea that the N-terminus is essential for prolonged interaction with nonspecific DNA. MSD analysis indicated a decrease in diffusivity with increasing DNA tension for both full-length and ΔN-term TDG, showing an ∼ 5-fold decrease from 5 to 40 pN (Fig. 4e-f, Table 2). Longer lifetimes and a decrease in diffusivity at higher tensions could indicate that TDG is better at interrogating the DNA for the base modification. Furthermore, these data indicate that bending of the DNA is not essential for TDG to bind productively to DNA. This idea is further supported by an increase in the residence time per base pair (Supplemental Note, Supplemental Table 2). The anomalous diffusion exponent, α, provides further insight into the movement of a particle, with α = 1 is random diffusion, α < 1 sub-diffusive (constrained motion), and α > 1 is super diffusive. For full-length WT TDG a decrease in the anomalous diffusion exponent, α, was observed but we did not see a significant decrease in α for ΔN-term TDG (Fig. 4e-f, Table 2). The TDG variants, N140A, R275L, and R275A, were also investigated at 40 pN to further investigate base interrogation investigate base interrogation at the idea of base interrogation. TDG N140A had a similar behavior to WT TDG: a decrease in both alpha and diffusivity were observed with an increase in tension (Fig. 4g). At high tension, the diffusivity of TDG R275L significantly decreased compared to low tension, but no significant difference was observed for TDG R275A. (Fig. 4h-i), supporting the idea that the size of the side chain aids in flipping in the base. These data indicate that DNA tension, perhaps through increasing the distance between stacked bases (∼5% increase in distance at 40 pN compared to 10 pN), provides better interaction of the N-terminal domain to increase the affinity to DNA increasing the dwell time and slowing the rate of diffusion to a mode of constrained motion.

**Table 2.** Binding lifetimes and diffusivities dependent upon tension.

| Protein | Tension (pN) | Lifetime(s)<sup>a</sup> | Diffusivity (μm<sup>2</sup>/s)<sup>b</sup> | α<sup>c</sup> |
| --- | --- | --- | --- | --- |
| Wild Type | 5 | 6.49 ± 0.04 | 0.034 ± 0.019 | 1.07 ± 0.22 |
|  | 10 | 6.45 ± 0.06 | 0.032 ± 0.027 | 1.05 ± 0.24 |
|  | 20 | 6.25 ± 0.20 | 0.026 ± 0.016 | 1.11 ± 0.27 |
|  | 30 | 10.31 ± 0.11 | 0.008 ± 0.006 | 0.83 ± 0.32 |
|  | 40 | 8.20 ± 0.27 | 0.007 ± 0.006 | 0.72 ± 0.35 |
| ΔN term | 5 | 0.99 ± 0.02 | 0.049 ± 0.067 | 0.99 ± 0.25 |
|  | 10 | 1.39 ± 0.02 | 0.050 ± 0.049 | 1.04 ± 0.35 |
|  | 20 | 1.58 ± 0.02 | 0.047 ± 0.052 | 0.98 ± 0.40 |
|  | 30 | 2.03 ± 0.02 | 0.020 ± 0.015 | 1.19 ± 0.46 |
|  | 40 | 2.03 ± 0.02 | 0.012 ± 0.009 | 0.99 ± 0.40 |
| N140A | 10 | 5.9 ± 0.32 | 0.025 ± 0.017 | 1.05 ± 0.29 |
|  | 40 | 5.52 ± 0.12 | 0.007 ± 0.005 | 0.85 ± 0.32 |
| R275A | 10 | 0.7 ± 0.03 | 0.239 ± 0.179 | 0.90 ± 0.43 |
|  | 40 | 0.73 ± 0.04 | 0.277 ± 0.403 | 1.01 ± 0.29 |
| R275L | 10 | 1.4 ± 0.37 | 0.143 ± 0.095 | 1.07 ± 0.34 |
|  | 40 | 0.66 ± 0.02 | 0.037 ± 0.031 | 1.05 ± 0.45 |

<sup>a</sup> The tau is the best fit value ± the standard error of the fit to the observed data
<sup>b</sup> The diffusivity is the mean ± the standard error of the mean
<sup>c</sup> The alpha is the mean ± the standard error of the mean

### N-terminus of TDG mediates nucleosome interactions

We sought to understand how chromatin structure impacts the TDG search mechanism by reconstituting an undamaged nucleosome using a 601 sequence^30^and ligating the nucleosome into 6.2 kb biotinylated handles using the DNA tethering kit from LUMICKS (Supplemental Fig. 7). We then performed single-imaging experiments using TDG-HaloTag and the Cy3 labeled mononucleosome-containing DNA substrate. We observed three different behaviors when TDG encountered the nucleosome: (1) TDG approaches from one side and hops over the nucleosome (bypass) (Fig. 5a), (2) TDG collides with but does not move past the nucleosome (no bypass) (Fig. 5b), and (3) TDG colocalizes with the nucleosome after the initial encounter (Fig. 5c). Further analysis revealed that TDG bypasses the nucleosome during 33.3% of encounters, did not bypass the nucleosome during 39.2% of encounters, and colocalized with the nucleosome during 27.5% of the encounters. CRTD analysis fit to a one-phase exponential indicated that TDG bound to nucleosomes with a lifetime of 16.5 ± 1.9 seconds (Fig. 5e), which is 2-fold longer than non-damaged DNA, and 4.5-fold shorter than binding to 5fC.

**Figure 5:**
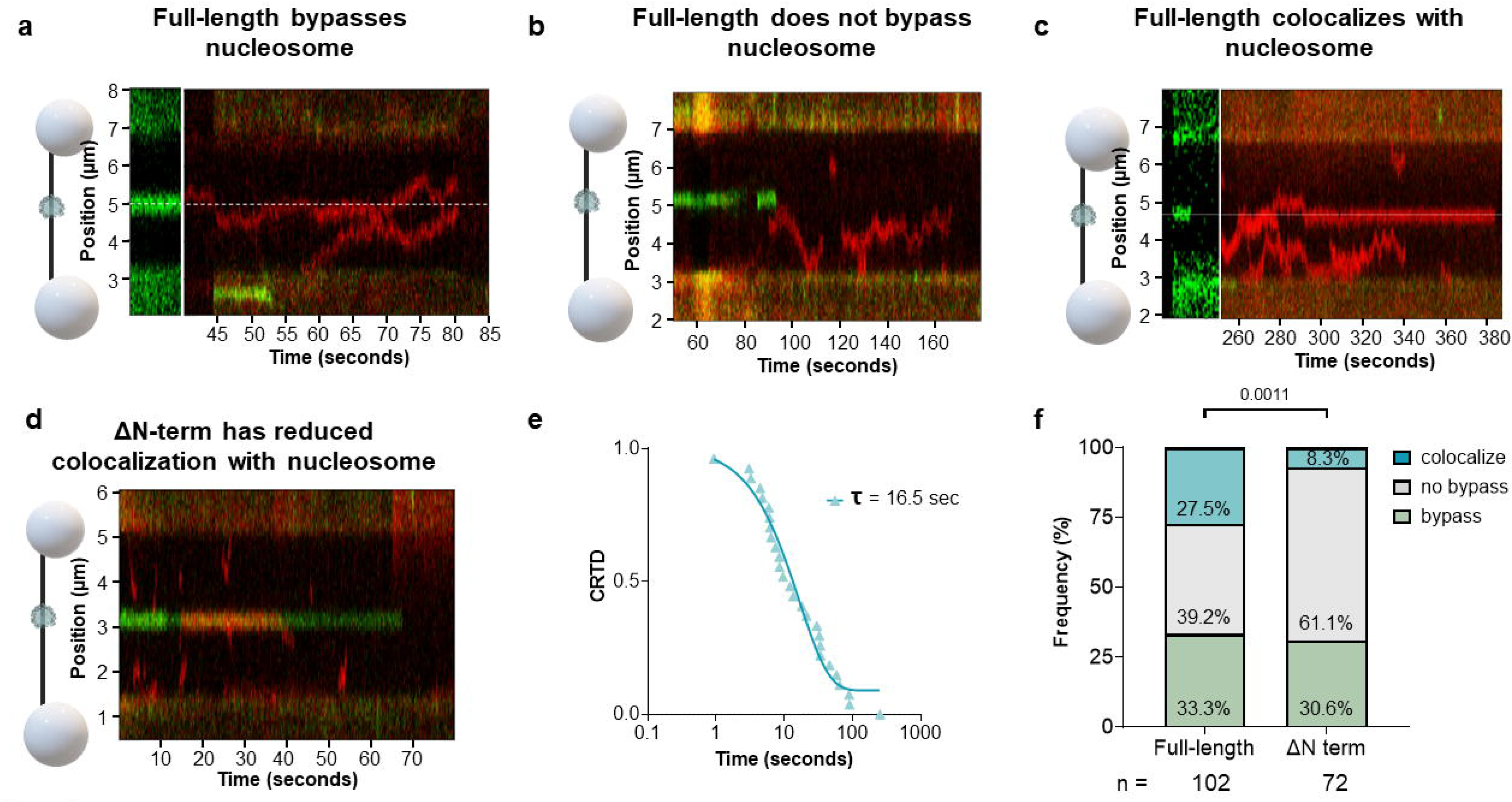
N-terminus of TDG is important for interacting with nucleosomes. A cartoon depiction of the DNA substrate with Cy3 labeled nucleosome. Representative kymograph with full-length TDG (shown in red) (a) bypassing, (b) does not bypass, or (c) colocalizes with the nucleosome shown in green. (d) Representative kymograph of ΔN-term TDG (red) colocalizing with nucleosome (green). (e) CRTD analysis fit to a one-phase decay of TDG interacting with the NCP (n = 28). (f) Stacked bar graph showing the fraction of bypass (green), no bypass (grey), and colocalized (teal) events for full-length and ΔN-term. shows there is *p* = 0.0011 by *Χ^2^*.

Since we observed the N-terminus is important for movement on DNA, we also sought to see if the N-terminus plays a role in binding or bypassing nucleosomes. To address this, we performed additional single-imaging experiments using TDG-HaloTag-ΔN-term and the mononucleosome-containing DNA substrate. These experiments revealed a significant decrease in the frequency of ΔN-term colocalized with the nucleosome (8.3%), an increase in the number of events that approached but did not bypass the nucleosome (61.1%), and saw a similar frequency for bypass (30.6%) (Fig. 5e). This decrease in TDG colocalization with the nucleosome indicates that the N-terminus plays an important role in nucleosome binding.

## Discussion

In this study, we utilized single-molecule methods to determine how TDG searches for DNA damage as well as uncovering how key active site residues, domains, and the presence of nucleosomes alter this search process. Similar to other DNA glycosylases, TDG forms moderately stable interactions with unmodified DNA to optimize its search^31,32^. Despite the ongoing 1D search, in our single molecule regime TDG primarily binds its modified sites directly out of solution in a 3D diffusion mechanism, with a limited number of events that bind unmodified DNA first and then slide into the lesion. Of note, sliding interactions of less than 100 base pairs are not discernable on our system so these events that appear as 3D may have some component of 1D diffusion not observed, as was observed for human uracil DNA glycosylase hUNG2^33,34^. Furthermore, we found that TDG bound specific lesions sites 10-fold longer than unmodified DNA. Since TDG is able to bind a wide array of modified bases, including 5caC and G:T mismatch, it is plausible that a similar behavior would be observed: TDG will bind to the modified base and become nonmotile, with rebinding to the generate abasic site. While it is currently not known how long TDG will bind to different lesions, it is likely that TDG will bind to a specific base modification for a time that is directly correlated to the biochemically determined cleavage rate followed by subsequent rebinding to an abasic site. Examining TDG under different ionic conditions, as well as two color experiment, demonstrated that TDG is able to both slide and hop along the DNA, which appears to be common amongst bacterial and mammalian glycosylases^33,35,36^. We estimated TDG’s search length on λ DNA utilizing the observed diffusion constant in 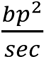 and the nonspecific lifetime of 7.9 sec (See Supplemental Note) to be 5225 bp or ∼10% of λ DNA per encounter.

The complex lesion search mechanism used by TDG has several potential benefits during interrogation of the genome for DNA lesions. Due to TDG’s high affinity for nondamaged DNA, the ability to switch between sliding and hopping may also be advantageous in situations where TDG encounters other factors on the DNA, including other transcription factors or nucleosomes (Fig. 6). With a sliding-exclusive method, full dissociation would need to occur to sample the DNA after the block. However, hopping would allow TDG to circumvent the block and continue scanning on the other side of an obstacle. This use of dual linear diffusion modes leads to the question of whether certain cellular conditions favor TDG hopping versus sliding on DNA. For example, if TDG has a sequence preference, it may adopt a conformation at these preferred sequences where tight binding leads to direct sliding as TDG further investigates that region of DNA. While it is currently unclear whether TDG has sequence preference, it is possible that sequences enriched in CG dinucleotides could result in initiating sliding behaviors. This type of increased binding to particular sequences would be highly advantages in regions with a heavier burden of DNA modification either due to (1) an enrichment of methylated CG dinucleotides or (2) TET enzymes localizing to methylated CpG rich islands to oxidize 5-methlyC groups for their removal by TDG and subsequent transcriptional activation^37^. It has been previously determined the glycosylase OGG1 slides and hops along the DNA in search of a lesion^31,35^, whereas hUNG2, a uracil DNA glycosylase family member, performs 3D diffusion, short range sliding, and hopping to find a lesion^33^. Thus, TDG exhibiting multiple modes of diffusion is in agreement with the behavior observed for other mammalian glycosylases and appears to be essential to efficiently scan the DNA and the ability to switch between modes may be dependent upon sequence or other cellular factors (Fig. 6a).

**Figure 6:**
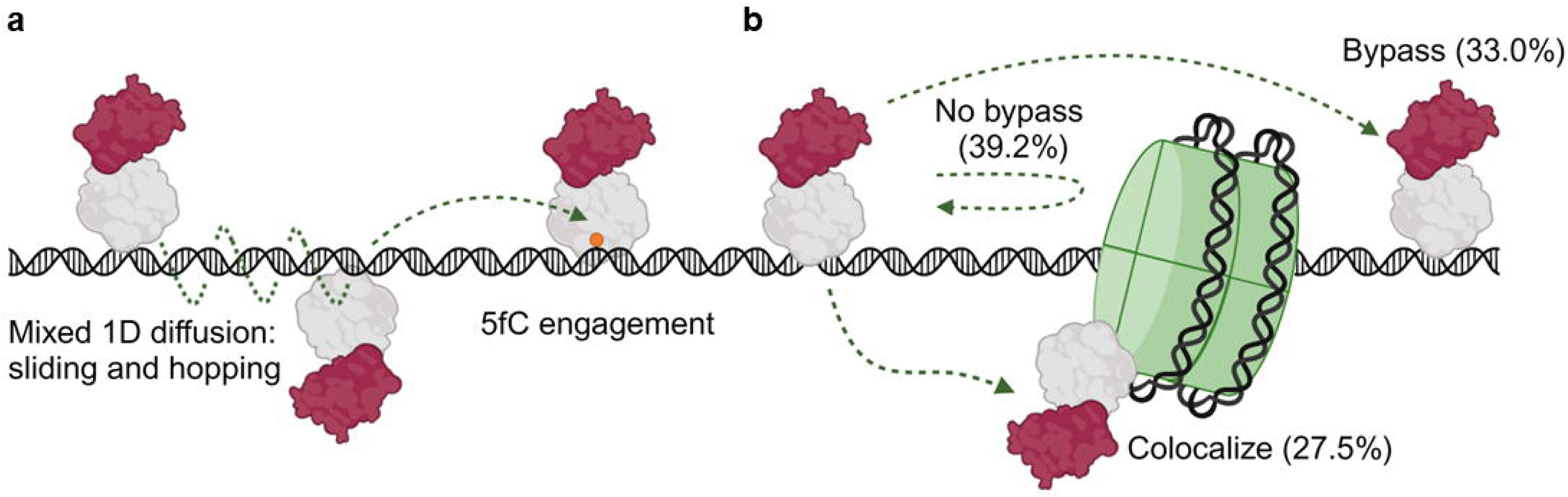
Working model. TDG uses multiple modes of linear diffusion to efficiently search for 5fC. (a) WT TDG (gray) fused to HaloTag (red) scans dsDNA with sliding and hopping interactions, with a lifetime of 7.9 ± 0.06 sec and diffusivity of 0.028 ± 0.02 µm2/sec. Upon 5fC engagement (orange), TDG binds with a lifetime of 72.9 ± 5.4 sec. These interactions are dependent on active site residues N140, R275, and the N-terminal domain. (b) When TDG encounters a nucleosome core particle, it can bypass, not bypass, or colocalize with the NCP (16.5 ± 1.9 sec). Importantly the N-terminal domain of TDG stabilizes the interaction with the NCP, resulting in a four-fold increase in colocalization frequency. Created with BioRender.com.

Our single molecule work also revealed the active site variants N140A, R275A, and R275L change the behavior of TDG on DNA. N140A had a shorter dwell time on 5fC but did not exhibit a faster diffusion than wild type. This contrasts other previous studies with the glycosylase OGG1, where the catalytically dead glycosylase bound much longer than that of the WT^19,38^. On the other hand, R275A/L variants both exhibited a shorter dwell time as well as a faster diffusion. One explanation for this behavior is that R275 probes the DNA sequence during the sliding, enabling efficient detection of DNA damage but also slowing the speed of sliding interactions. Further, it is possible that the lack of the positive charge causes TDG to have lower binding affinity and thus shorter dwell times and faster rates of diffusion. A previous study on bacterial glycosylases Fpg, Nei, and Nth has also shown that if a residue that interrogates the DNA is changed to alanine, faster diffusivity was observed, indicating that efficient recognition of a base is dependent on a single active site residue^27^. Together, these findings provide an in-depth description of how key catalytic residues of TDG contribute to its search for DNA damage and provides insight into how its mixed mode of hopping and sliding searches contribute to efficient damage detection and repair.

Deleting the disordered N-terminus of TDG, consisting of the first 81 amino acids, did not reduce the ability to bind to 5fC. However, the ability of ΔN-term TDG to search nonspecific DNA was significantly reduced, due to the three-fold shorter lifetime of ΔN-term TDG bound to DNA nonspecifically. While the tension on DNA at any given point in the cell is not well known, it is known that nucleosomes begin to unwrap at 5 pN^39^, hairpins begin to pull apart 10-15 pN^40,41^, G-quadruplexes begin to unfold 15-20 pN^42^, RNA polymerase has been shown to exert a force of 25-30 pN on DNA^43^ and double strand DNA begins to unwind at ∼50 pN^44,45^. With increasing tension, the λ DNA may begin to have regions that may potentially allow for TDG to interrogate the DNA more easily perhaps due to widening of the base steps and less stacking interactions, allowing better interrogation of the DNA at higher tensions. The observed decrease in α, indicating more constrained motion on DNA, for full-length but not ΔN-term TDG suggests that the disordered N-terminus may play a role in allowing TDG to thoroughly interrogate the DNA for a lesion. The N-terminus may also play a role in TDG switching between hopping and sliding. If TDG needs to form for tighter interactions with the DNA in order to slide, the N-terminus, which contains several positively charged amino acid residues, may adopt a confirmation that limits hoping to better probe an area. Similarly, a study has shown that the N-terminus of hUNG2 has an increase in contact with DNA during molecular crowding which results in an efficient sliding and prevents hUNG2 from dissociating and diffusing into bulk solution^46^.

We have shown that not only is TDG able to hop over a nucleosome, but it also interacts with a nucleosome with an apparent dwell time of 16.5 ± 1.9 seconds. Deleting the N-terminus significantly reduces the binding frequency to nucleosomes (Fig. 6b). This interaction of TDG with nucleosomes may act as a scaffold to help facilitate removal of oxidized cytosine in heterochromatic regions. Alternatively, perhaps TDG associates with nucleosomes prior to the arrival of TET enzymes, and thus would increase the efficiency of oxidative demethylation by already being in the correct location prior to the initial 5mC oxidation step in heterochromatic regions. Efficient active oxidative demethylation is vital for life and of interest for potential therapeutic applications^47^, and this work gives key mechanistic insight on this process. TDG interacting with a nucleosome could also alter nucleosome dynamics and may interfere with fork progression and DNA replication, and thus might explain its degradation during S-phase in mammalian cells.

Taken together, our studies offer new insight into how key active site residues and domains of TDG contribute to its efficient search for DNA damage, and how chromatin can act either as a roadblock or a scaffold during the search process. Of note, these studies with nucleosome core particles represent a unique look at how base excision repair enzyme interact with nucleosomes that do not have free DNA ends present. Future studies will need to be completed to determine the impact nucleosome unwrapping, chromatin remodelers, and how post translational modifications of TDG will impact its ability to efficiently search for its substrates.

## Online Methods

### DNA substrate preparation

#### Nick translation for confocal imaging

Lambda DNA was biotinylated as described previously^19^. Briefly, DNA was purchased from New England Biotechnologies and treated with Klenow fragment polymerase (NEB) and a dNTP mix containing biotinylated dCTP. This results in one side of the DNA containing four biotins and the other containing six biotins.

Next, 1 µg of the biotinylated lambda DNA is treated with Nt.BspQI (NEB) for 1 hour at 50°C then heat inactivated for 20 minutes at 80°C, which will generate single strand breaks at 10 different sites along the DNA by cutting 3’ of the recognition site 5’-GCTCTTCN-3’. To incorporate 5fC at the 10 sites, 800 ng nicked DNA was incubated with a dNTP mix containing d5fCTP, fluorescein labeled dUTP, dATP, and dGTP were incubated and 10 units of DNA polymerase 1 (NEB) for 6 minutes at 37°C. Reaction was inactivated by heating at 75°C for 20 minutes and allowed to slowly cool to room temperature. To seal the nicks, a reaction containing 800 ng of the nick translated DNA, 8 mM ATP and 10 units of T4 ligase (NEB) were incubated overnight at 16°C. Reaction was heat inactivated for 10 minutes at 65°C and allowed to cool to room temperature slowly. Of the 10 d5fCTP sites, only eight d5fCTP sites can be resolved as two of the sites are within 400 bp of each other and one site is near the end of λ DNA and thus too close to the bead (Supplemental Fig. 4a-c).

#### Arms ligation for confocal imaging

Using the LUMICKS DNA tethering kit and protocol, a sequence of interest is ligated into two handles that are each 6.3 kb in length to generate a substrate that is 12.6 kb in total. For the defined 5fC, the following oligonucleotide sequences were ordered from Trilink and IDT, respectively:

Top strand (5fC): 5’-phosphate-*CAAC* ACC AGT CCA TCG CTC A**5fC**G TAC AGA GCT G–3’ Bottom strand: 5’-phosphate – *ACCA* CAG CTC TGT ACG TGA GCG ATG GAC TGG T-3’ Annealing reactions were heated at 95°C for 5 minutes in buffer containing 10 mM Tris-HCl (pH 8.0) and 100 mM KCl, and the heat block was turned off and reactions then cooled slowly to room temperature. In a 10 µL reaction, 25 nM of the duplex, 2.5 µL of each handle, 1 µL of 10X ligase buffer, and 0.5 µL of ligase were mixed and incubated at 16°C overnight. Reaction was heat inactivated at 65°C for 10 minutes and allowed to slowly cool to room temperature. Substrate was stored at 4°C and protected from light.

Undamaged nucleosome were reconstituted on a 601 sequence as described previously^30^. Briefly, Human histone H3 C96S C110A, H2A K119C, H2B, and H4 were ordered from the Histone Source at Colorado State University. Incubate H2A K119C and H2B in 2 mg/mL guanidinium buffer at room temperature for 2 hours. Mix equimolar amounts of H2A and H2B and dialyze against high salt refolding buffer a total of 3 times, and at least 8 hours for each exchange at 4°C. Repeat the same process for H3 C96S C110A and H4 to refold the tetramer. Purify the H2A K119C/H2B dimer and the H3 C96S C110A/H4 tetramer over Superdex 200 column. Incubate the H2A K119C/H2B dimer with 0.7mM TCEP for 20 minutes at 4°C. Cy3-maleimide dye was added to the H2A K119C in a 2:1 molar ratio and incubated at room temperature for 1 hour while rocking. Reaction was quenched with 10 mM DTT, the dimer was purified over a Superdex S200 column, and frozen in 50% glycerol. After confirming the stoichiometry with SDS gel, add equal volume of 100% glycerol to store the H2A K119C/H2B dimer and the H3 C96S C110A/H4 tetramer at -20°C.

To reconstitute the nucleosome on DNA, the following ultramer sequences were ordered from IDT:

Top Strand: 5’-phosphate-*CAAC* TGA GAC CAT GTA CCC AGT TCG AAT CGG ATG TAT ATA TCT GAC ACG TGC CTG GAG ACT AGG GAG TAA TCC CCT TGG CGG TTA AAA CGC GGG GGA CAG CGC GTA CGT GCG TTT AAG CGG TGC TAG AGC TGT CTA CGA CCA ATT GAG CGG CCT CGG CAC CGG GAT TCT CGA TAA CTC AGC AAT AGT GGG TCT CA – 3’

Bottom strand: 5’-phosphate -*ACCA* TGA GAC CCA CTA TTG CTG AGT TAT CGA GAA TCC CGG TGC CGA GGC CGC TCA ATT GGT CGT AGA CAG CTC TAG CAC CGC TTA AAC GCA CGT ACG CGC TGT CCC CCG CGT TTT AAC CGC CAA GGG GAT TAC TCC CTA GTC TCC AGG CAC GTG TCA GAT ATA TAC ATC CGA TTC GAA CTG GGT ACA TGG TCT CA – 3’

The annealed DNA, H2A K119C/H2B dimer, and the H3 C96S C110A/H4 tetramer are mixed in a 1:2:1 molar ratio and equilibrate in dialysis tubing against high salt buffer for 30 minutes. To remove the high salt buffer, a series of dialysis steps occur to transition from 1.5 M NaCl to 0.125 M NaCl. The reconstituted nucleosome is concentrated, and heat shocked at 55°C for 30 minutes, prior to spinning over a 10-40% sucrose gradient for 40 hours at 125,000 xG at 4°C. Fractions containing reconstituted nucleosomes are combined, buffered exchanged into TE buffer, and concentrated to ∼1 µM and stored at 4°C. To ligate into the LUMICKS handles, 25 nM of the reconstitute nucleosome, 2.5 µL of each handle, 1 µL of 10X ligase buffer, and 0.5 µL of ligase were mixed and incubated at 16°C overnight. Substrate was stored at 4°C and protected from light.

### Plasmid Constructs

Wild type TDG-HaloTag and TDG-HaloTag variants were expressed from plasmids constructed utilizing Gene Universal Inc with the pHTN HaloTag CMV-neo vector (Promega). (Supplemental Figs. 1a-e)

### Nuclear Extract

Nuclear extracts were prepared as described previously^19^. Briefly, U2OS cells were cultured in 5% CO2 in Dulbecco’s modified Eagle’s medium (DMEM) supplemented with 4.5 g/L glucose, 10% fetal bovine serum (Gibco), and 5% penicillin/streptavidin (Life Technologies). To obtain transient overexpression of TDG-HaloTag, cells were transfected with 4 µg of plasmid per 4 million cells using the lipofectamine 2000 reagent and protocol (Thermo Fisher Cat# 11668019). Nuclear extraction was performed 24 hours after transfection utilizing nuclear extraction kit from Abcam (ab113474). Following the protocol from the kit, 10 µL single-use aliquots were made, flash-frozen and, stored at -80°C. Extracts were diluted in buffer containing 20 mM HEPES pH 7.5, 100 mM NaCl, 0.2 mM EDTA, 0.1 mg/ml BSA, 1 mM DTT, 1 mM Trolox, and 5% glycerol.

### Nuclear Extract Quantification

#### SDS gel

A total of 2.5 µg of each nuclear extract was loaded in duplicate on a 4-12% Bis Tris gels (Invitrogen NP0323BOX) with a dye free sample buffer (4X buffer: 40% glycerol, 200 mM Tris-Cl (pH 6.8), and 8% SDS). The gel was imaged using a laser scanner for Cy5 (Typhoon, Amersham). Gels were then stained with Coomassie blue and imaged. The total intensity for each lane was measured and the lanes were averaged together in order to normalize for loading. Finally, each of the variant extracts were normalized to the wild type TDG in order to look at expression levels and the level of free dye. (Supplemental Figure 2a-c)

#### Western Blotting

Different amounts of purified proteins and nuclear extracts were denatured at 95°C for 10 minutes. Equal volumes were loaded on a 4-20% tris-glycine polyacrylamide gel (Invitrogen XP04202BOX). Proteins were transferred onto a polyvinylidene difluoride membrane and blocked in 20% nonfat dry milk in PBST (1x phosphate-buffered saline containing 0.1% Tween 20) for 1 hr at room temperature. Membranes were probed with TDG primary antibody (1:500 Sigma HPA052263) overnight at 4°C. Membranes were washed 3 times for 14 min in PSBT and incubated with peroxidase conjugated rabbit secondary antibody for 1 h at room temperature. Membranes were washed again 3 times for 15 minutes before developing using SuperSignal West Femto Maximum Sensitivity Substrate (Thermo Fisher Scientific #34095). (Supplemental Figure 2d),

### Confocal Imaging in the LUMICKS C-trap

#### Imaging

Fluorophores and dyes utilized were excited with the laser closest to their maximum intensity. Fluorescein was excited with a 488 nm laser and emission collected with a 500–550 nm band pass filter, Cy3 and HaloTag-JF552 were excited with a 561 nm laser and the emission was collected with a 575–625 nm band pass filter, and HaloTag-JF635 was excited with a 638 nm laser and emission collected with a 650– 750 nm band pass filter. All data were collected with a 1.2 NA 60x water emersion objective and photons measured with single-photon avalanche photodiode detectors. The laser power was set to 5% and kymographs were collected continuously with 0.1 msec exposure for each pixel of size 100 nm, which resulted in approximately 30 frames/sec for λ DNA. All data was collected with a minimum of two different preparations of nuclear extracts across multiple days.

Single-molecule experiments were then performed with a LUMICKS C-trap using a 5-channel flow chamber. For experiments using λ or nick translated DNA, ∼4 µm streptavidin coated beads and experiments using the LUMICKS handles, ∼1.7 µm streptavidin coated beads were used. The beads were captured using the dual optical tweezers and then transferred to a second chamber containing the biotinylated DNA and the biotinylated DNA molecule tethered between the streptavidin coated beads. Once the DNA molecule is tethered, it is transferred to a final chamber containing a 1:10 dilution of the nuclear extract with TDG-HaloTag-JF635. To determine the concentration of the protein in the flow cell, background photon intensities were determined to generate the standard curve of purified HaloTag labeled with JF635 or JF552. The background intensities of the protein from the nuclear extract were measured and the curve was utilized to determine protein concertation (Supplemental Figure 3a-b).

#### Analysis

Kymographs were analyzed with custom software from LUMICKS, Pylake. The kymographs were exported as .h5 files from the C-trap and viewed using the utility “C-Trap .h5 Visualization GUI (2020)” by John Watters (harbor.lumicks.com). Line tracking was performed using a custom script from LUMICKS based on a Gaussian fit over the line to determine its positions^48^. Only events that associated and dissociated during the course of a 5-to-10-minute kymograph were tracked. Events are plotted on cumulative residence time distribution (CRTD) plot in order to determine the binding lifetime. Specific binding event curves are fit with a two-phase exponential decay, for catalytically active WT protein. One-phase exponential decays were sued for the variants, which have little or no catalytic activity, and nonspecific binding events for the variants and WT. The two-phase fit for WT represents a second rebinding event to the catalytic product abasic site with a shorter dwell time. The goodness of the fit for the CRTD curves range from *R^2^* = 0.96 – 0.99. Mean squared displacement (MSD) analysis was performed using a custom script from LUMICKS based on the following equation:

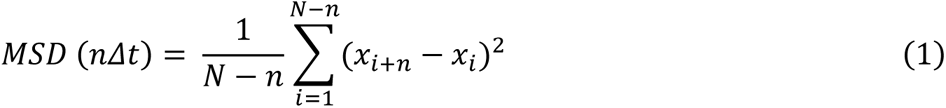

Where *N* is the total number of frames in the phase, *n* is the number of frames at a given time step, *Δt* is the time increment of one frame, and *x_i_* is the particle position in the *i*th frame^27^. The diffusion coefficient (*D*) was determined by fitting a linear model of one-dimensional diffusion to the MSD plots:

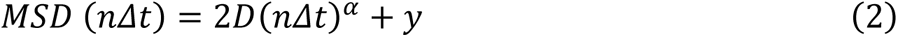

where *α* is the anomalous diffusion coefficient and *y* is a constant (y-intercept). Fittings resulting in *R^2^* < 0.8 or using <10% of the MSD plot were not considered.

## Supporting information

Supplemental Math

## Data Availability

All data are available from the corresponding author upon reasonable request.

## Code Availability

Python scripts used to calculate lifetime and MSD analysis have been deposited at https://harbor.lumicks.com/scripts.

## Contributions

B.L.S. and B.V.H. conceived the research. B.L.S. performed all Ctrap and protein concentration determination experiments. V.R. performed all western blot experiments. B.L.S analyzed all single-molecule data. T.M.W. reconstituted nucleosomes. B.L.S., M.A.S. and B.V.H. drafted the paper, which was reviewed, discussed, and edited by all authors.

## Acknowledgements

We greatly appreciate the discussions with the lab members during the course of these experiments. This work was supported by NIH R35ES031638 (BVH), T32GM088119 (BLS), the Hillman Postdoctoral Fellowship for Innovative Cancer Research and F32ES034982 (MAS), 2P30CA047904 to the UPMC Hillman Cancer Center, the major equipment grant S10OD032158-01A1 (BVH), R35-GM136225 (ACD), R35GM128652 (BDF), and F32GM140718 (TMW).

