## Supplemental Math for "Thymine DNA glycosylase combines sliding, hopping, and nucleosome interactions to efficiently search for 5-formylcytosine"

### Supplemental Note

#### Estimation of the Hydrodynamic Radii

The calculations to determine the hydrodynamic radii were followed as described previously [1-3]. The radius of gyration for full length TDG was estimated by importing the AlphaFold structure in to WinHYDROPRO [4]. Additionally, PDB 6U32 was imported to determine the radius of gyration for HaloTag. The following parameters were utilized with the "shell model from atomic level" option to estimate the radius: MW = 48 kDa (HaloTag) and MW = 34.28 kDa (HaloTag), room temperature T = 20°C, and viscosity of water at room temperature $\eta = 0.89 \times 10^{-3}$ Pa · s.

$$R_H = \frac{R_g}{\rho} \tag{1}$$

$$R_{H,TDG} = \frac{R_{g,TDG}}{\rho} = \frac{3.69\ nm}{\left(\frac{3}{5}\right)^{\frac{1}{2}}} = 4.76\ nm$$

$$R_{H,HaloTag} = \frac{R_{g,HaloTag}}{\rho} = \frac{1.85\ nm}{\left(\frac{3}{5}\right)^{1/2}} = 2.39\ nm$$

Where ρ is the ratio of the geometry to the hydrodynamic ratio for a spherical molecule [5].

To determine the effective hydrodynamic radii of HaloTag fused to TDG:

$$R_{H,eff} = \left({R_{H,TDG}}^3 + {R_{H,HaloTag}}^3\right)^{\frac{1}{3}} \tag{2}$$

$$R_{H,eff} = (4.76^3\ nm^3 + 2.93^3\ nm^3)^{1/3} = 4.95\ nm$$

#### Estimation of Theoretical Limit of Diffusion Coefficient

The diffusion coefficient can be described using the Stokes-Einstein equation:

$$D = \frac{k_B T}{\xi} \tag{3}$$

where ξ is the friction term and $k_B T$ at room temperature is $4.11 \times 10^{-21}$ J. The friction term for a protein sliding on the DNA following the helical axis was determined by Schurr [6] and modified by Bagchi [7] to be as follows:

$$\xi = 6\pi\eta R_{H,eff} + \left(\frac{2\pi}{10.5\ BP}\right)^2 \left[8\pi\eta R_{H,eff}^3 + 6\pi\eta R_{OC} R_{H,eff}^2\right] \tag{4}$$

where $BP$ is the distance between two base pairs (0.34 nm), viscosity of water at room temperature $\eta = 0.89 \times 10^{-3}$ Pa · s and $R_{OC}$ is the occ-center distance from the center of mass of the protein to the helical axis of the DNA.

$$R_{OC} = R_{H,eff} + 1\ nm \tag{5}$$

$$R_{OC} = 4.95\ nm + 1\ nm = 5.95\ nm$$

Combining the friction term with the Stokes-Einstein equation allows for the calculation of the theoretical limit of diffusion of the HaloTag fused to TDG sliding along DNA:

$$D_{lim} = \frac{k_B T}{6\pi\eta R_{H,eff} + \left(\frac{2\pi}{10.5\ BP}\right)^2 \left[8\pi\eta R_{H,eff}^3 + 6\pi\eta R_{OC} R_{H,eff}^2\right]} \tag{6}$$

$$D_{lim} = 0.256\ {}^{\mu m^2}/_{s}$$

**<u>Calculating the energy barrier to free diffusion:</u>**

$$k = e^{\frac{-E_A}{k_B T}} \tag{7}$$

where $k$ is the rate constant in *steps/s* (in this instance it is the stepping rate $\frac{2D}{BP^2}$, where $BP^2$ is assumed to be 1). The energy barrier to the free diffusion is the difference between the theoretical (aka the "barrier-less") $E_A$ and the experimentally determined $E_A$. By rearranging the equation to solve to for $E_A$

$$\ln k = \frac{-E_A}{k_B T} \tag{8}$$

$$-E_A = \ln k \;\, x \; k_B T$$

$$\Delta E_A = [\ln\left(\frac{2D_{lim}}{BP^2}\right) - \ln\left(\frac{2D_{expt}}{BP^2}\right)] \;\; x \; k_B T$$

$$\Delta E_A = \ln\left(\frac{\frac{2D_{lim}}{BP^2}}{\frac{2D_{expt}}{BP}^2}\right) x \; k_B T$$

$$\Delta E_A = \ln\left(\frac{D_{lim}}{D_{expt}}\right) x \; k_B T \tag{9}$$

Using the equation, the energy barrier is $2.77 \pm 0.75 \; k_B T$ for WT TDG in 75 mM NaCl, $2.21 \pm 0.75 \; k_B T$ for WT TDG in 100 mM NaCl, and $2.11 \pm 0.71 \; k_B T$ for WT TDG in 150 mM NaCl. The energy barrier for the variants was calculated to be $2.45 \pm 0.82 \; k_B T$ for N140A, $0.68 \pm 0.65 \; k_B T$ for R275A, and $1.08 \pm 0.8 \; k_B T$ for R275L.

For the tension series the, the energy barrier for full-length TDG is $2.02 \pm 0.56 \, k_B T$ for 5 pN, $2.08 \pm 0.84 \, k_B T$ for 10 pN, $2.29 \pm 0.62 \, k_B T$ for 20 pN, $3.47 \pm 0.75 \, k_B T$ for 30 pN, and $3.6 \pm 0.86 \, k_B T$ for 40 pN. The energy barrier for ΔN term TDG is $1.98 \pm 1.37 \, k_B T$ for 5 pN, $1.96 \pm 0.98 \, k_B T$ for 10 pN, $2.02 \pm 1.11 \, k_B T$ for 20 pN, $2.87 \pm 0.75 \, k_B T$ for 30 pN, and $3.68 \pm 0.56 \, k_B T$ for 40 pN.

**Estimation of residence time at each base pair:**

The stepping rate can be utilized to determine the dwell time ($\tau_{bp}$) TDG-HaloTag spends on one base pair:

$$\tau_{bp} = \frac{1}{k} = \frac{1}{2D/_{l^2}} = \frac{l^2}{2D} \tag{10}$$

Where $D$ is the experimentally determined diffusivity, and $l$ is assumed to be 1 base pair. The residence time was estimated to be $3.6 \pm 2.7$ μ$s$ for wild type in 75 mM NaCl, $2.1 \pm 1.6$ μ$s$ for wild type in 100 mM NaCl, $1.9 \pm 1.3$ μ$s$ for wild type in 150 mM NaCl, $2.6 \pm 2.2$ μ$s$ for N140A, $0.5 \pm 0.3$ μ$s$ for R275A, and $0.7 \pm 0.5$ μ$s$ for R275L.

For the tension series the, the residence time for full-length TDG was estimated to be $1.7 \pm 1.0$ μ$s$ for 5 pN, $1.8 \pm 1.5$ μ$s$ for 10 pN, $2.3 \pm 1.4$ μ$s$ for 20 pN, $7.3 \pm 5.5$ μ$s$ for 30 pN, and $8.3 \pm 7.1$ μ$s$ for 40 pN. The residence time for ΔN term TDG was estimated to be $1.2 \pm 1.6$ μ$s$ for 5 pN, $1.2 \pm 1.1$ μ$s$ for 10 pN, $1.2 \pm 1.4$ μ$s$ for 20 pN, $2.9 \pm 2.2$ μ$s$ for 30 pN, and $4.9 \pm 3.6$ μ$s$ for 40 pN.

**Calculating the expected sliding length:**

In a previous study, they derived an equation dependent upon the diffusivity and mean lifetime of binding events to determine how far a protein is capable of sliding on the DNA[8]

$$expected\ sliding\ length = \sqrt{\frac{\pi * D_{expt}}{2 * \lambda}} \tag{11}$$

Where 1/λ is the mean lifetime and $D_{expt}$ is the diffusivity in $bp^2$/sec

$$expected\ sliding\ length = \ 5224.98\ bp$$

The full length of λ DNA is 48.5 kb. TDG is capable of searching 5225 bp, or 10% of λ DNA in 7.9 seconds.
